# Hippocampal purinergic P2X7 receptor level is increased in Alzheimer’s disease patients, and associated with amyloid and tau pathologies

**DOI:** 10.1101/2024.02.28.582443

**Authors:** Cinzia A. Maschio, Junlong Wang, Upasana Maheshwari, Annika Keller, Axel Rominger, Uwe Konietzko, Agneta Nordberg, Christoph Hock, Roger M. Nitsch, Ruiqing Ni

## Abstract

**INTRODUCTION:** The purinergic receptor P2X7R, which is expressed on microglia and astrocytes, plays an important role in Alzheimer’s disease (AD). We aimed to characterize the alterations in P2X7R expression in AD patients by APOE ε4 allele, age and sex, as well as its association with amyloid and tau pathology.

**METHODS:** P2X7R staining and quantitative analysis of amyloid, tau, astrocytes and microglia were performed on postmortem hippocampal tissues from 35 AD patients; 31 nondemented controls; caudate/putamen tissue from corticobasal degeneration (CBD), progressive supranuclear palsy (PSP) patients; and bran tissue from aged 3×Tg mouse model of AD.

**RESULTS:** Activated microglia and reactive astrocytes were observed in the hippocampi of AD patients and exhibited altered morphology with denser cells and pronounced ramifications. Hippocampal P2X7R intensity was greater in the hippocampal subfields of AD patients than in those of nondemented controls and was correlated with amyloid level and Braak stage and was not affected by sex, APOEε4 allele, or age. P2X7R expression increased around Aβ plaques, cerebral amyloid angiopathy, tau inclusions in the hippocampus from AD patients and tau inclusions in the caudate/putamen from CBD and PSP patients.

**DISCUSSION:** We found an increased hippocampal P2X7R level in AD compared to non-demented control, which correlated with amyloid and tau pathologies. P2X7R is a potential marker for neuroinflammation in AD.

## Introduction

Alzheimer’s disease (AD) is the most common cause of dementia and is pathologically characterized by amyloid-beta (Aβ) plaque and tau tangle deposition (64). Primary tauopathies such as frontotemporal lobar degeneration (FTLD), corticobasal degeneration (CBD) and progressive supranuclear palsy (PSP) are pathologically characterized by disease-specific tau inclusions. Neuroinflammation is considered to play a pivotal role in the pathophysiology of AD. Resident microglia and astrocytes are the main components of the innate immune system and play important roles in AD and primary tauopathies (11, 26, 27). Genome wide association studies and single-cell genomic studies in AD have identified many risk genes that expressed in microglia and the critical role of neuroimmune pathways chronic inflammatory states for AD (38, 55). Moreover transcriptomic studies have found diverse and substantial astrocyte and microglial responses to show the most changes in response to AD pathology in murine and human studies (34, 53, 63). Microglial activation visualized by positron emission tomography (PET) has been shown to correlate with Aβ and tau loads (12, 14, 71), propagate jointly with tau across Braak stages (49), and cognitive decline in AD (41). The plasma level of glial fibrillary acidic protein (GFAP), a marker of astrocyte activation, has been shown to reflect the accumulation of Aβ, the presence of neurofibrillary tangles, and cognitive status (6, 30, 50) and can also predict the progression of AD (6, 15, 23).

Purinergic signalling mediated by purinergic receptors, particularly P2X7Rs, plays important roles in chronic immune and inflammatory responses (5, 18, 56). P2X7R is an adenosine triphosphate (ATP)-gated ion channel expressed on astrocytes (5), microglia and oligodendrocytes in the brain and macrophages in the periphery. P2X7R is expressed at low levels under physiological conditions when the extracellular ATP concentration is below the threshold required for activation, while it is activated when ATP levels rise above the threshold under pathological conditions. Notably, P2X7R has proinflammatory functions involving the activation of the NOD-, LRR- and pyrin domain-containing protein 3 (NLRP3) inflammasome; the secretion of proinflammatory cytokines; and the maturation of interleukin-1β (40, 54, 60). A possible protective role against AD has been reported for the 489C>T *p2rx7* gene polymorphism (61). In AD, P2X7R is involved in multiple pathological processes, including oxidative stress, and chronic neuroinflammation. An association between P2X7R and Aβ and tau has been reported: Aβ blunts the adenosine A2A receptor-mediated control of the interplay between P2X7R and P2Y1R-mediated calcium responses in astrocytes (19). P2X7R plays a pivotal role in promoting proinflammatory pathways via Aβ-mediated chemokine release (42). P2X7R inhibition has been shown to reduce amyloid plaques in animal models of AD by regulating the ability of microglia to phagocytose Aβ and via glycogen synthase kinase-3β and by influencing alpha-secretase-dependent processing (20, 43). P2X7R inhibition ameliorates ubiquitin□proteasome system dysfunction associated with AD (7) and reduces the release of tau-containing exosomes and tau-induced toxicity (17, 59); deficiency of the *p2rx7* gene has demonstrated efficacy in enhancing plasticity and cognitive abilities in tauopathy mice and amyloidosis mice (13, 42). One study showed that P2X7R regulate neuroinflammation in aged rat model by increasing Interleukin-1 alpha and Interleukin-1 beta release and suppressing the release of chemokine ligands, and Interleukin 6 (52). Moreover, P2X7R influences the tau inclusion burden in human tauopathies such as FTLD and induces distinct signalling pathways in microglia and astrocytes (4). This evidence has implicated P2X7Rs as therapuetic and biomarker targets for AD (68).

Here, we aimed to assess the alterations in P2X7R expression in postmortem hippocampi from NCs and AD patients by immunofluorescence staining, and understand the associations between P2X7R with tau pathology, Braak stage, and Aβ pathology, and influenced by age, apolipoprotein E (APOE) ε4 and sex.

## Materials and Methods

### Postmortem human brain tissue

Thirty-five AD patients, one PSP patient, one CBD patient, each with a clinical diagnosis confirmed by pathological examination, and thirty-one NCs were included in this study (detailed information in **Table 1**). Paraffin-embedded autopsy hippocampus tissue blocks were obtained from the Netherlands Brain Bank (NBB), Netherlands. All materials were collected from donors or from whom written informed consent was obtained for a brain autopsy, and the use of the materials and clinical information for research purposes were obtained by the NBB. The study was conducted according to the principles of the Declaration of Helsinki and subsequent revisions. All the autopsied human brain tissue experiments were carried out in accordance with ethical permission obtained from the regional human ethics committee and the medical ethics committee of the VU Medical Center for NBB tissue. Information on the neuropathological diagnosis of AD (possible, probable, or definite AD) or not AD was obtained from the NBB. Information on the Consortium to Establish a Registry for AD (CERAD), which applied semiquantitative estimates of neuritic plaque density and the Braak score based on the presence of neurofibrillary tangles (NFTs), is provided in Table 1. Patients with pathology other than AD were excluded from the study.

**Table 1.**
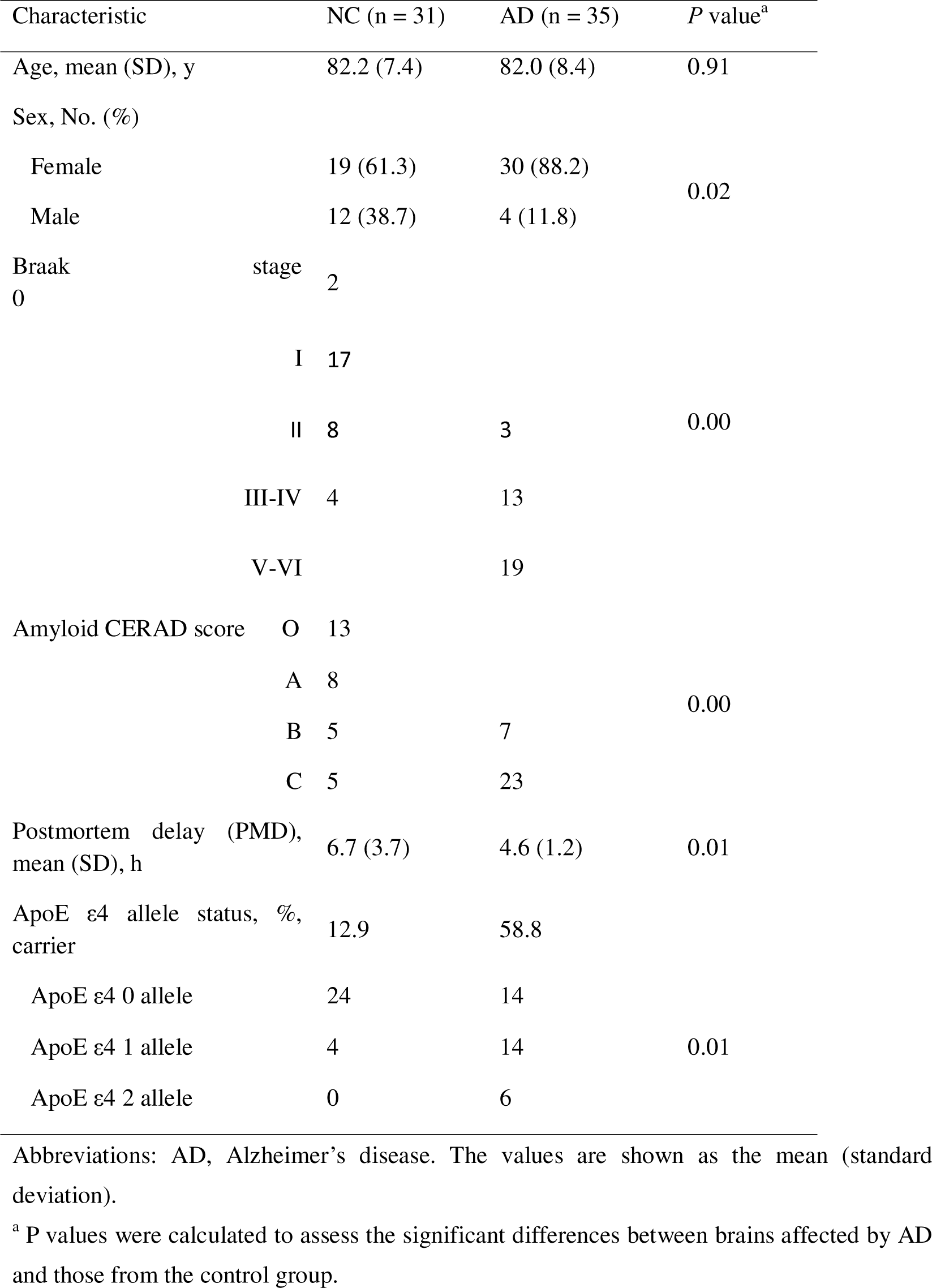
Demographic and neuropathologic characteristics of the 64 AD and NC postmortem human brains analysed.

### Animal models

Two 27-month-old 3×Tg mice [B6;129-Psen1tm1MpmTg (APPSwe, tauP301L)1Lfa/Mmjax] from the Jackson Laboratory (USA) and two age-matched wild-type mice were used (36, 47). Animals were housed in ventilated cages inside a temperature-controlled room under a 12-h dark/light cycle. Pelleted food (3437PXL15, CARGILL) and water were provided ad libitum. Paper tissue and red Tecniplast Mouse House® (Tecniplast, Milan, Italy) shelters were placed in cages for environmental enrichment. All the experiments were performed in accordance with the Swiss Federal Act on Animal Protection and were approved by the Cantonal Veterinary Office Zurich. All the experiments followed the ARRIVAL 2.0 guidelines. 3×Tg mice and wild-type mice were perfused under ketamine/xylazine/acepromazine maleate anaesthesia (75/10/2 mg/kg body weight, i.p. bolus injection) with ice-cold 0.1 M phosphate-buffered saline (PBS, pH 7.4) and 4% paraformaldehyde in 0.1 M PBS (pH 7.4), fixed for 24 h in 4% paraformaldehyde (pH 7.4) and then stored in 0.1 M PBS (pH 7.4) at 4°C.

### Immunohistochemistry and immunofluorescence staining

Paraffin-embedded fixed postmortem human brain tissues (from 34 AD patients and 30 NCs) were cut into 3 µm sections using a Leica microtome (Leica Microsystems, Germany) (46, 67). Hematoxylin and eosin (H&E) staining was performed according to routine procedures for each patient to provide anatomical information and to determine whether there were abnormalities in the brain. 4G8 (Aβ17-24), GFAP and Iba1 immunohistochemical staining and P2X7R immunofluorescence staining were performed on 34 AD and 30 NC postmortem human brain tissues to visualize the morphology and pattern of the hippocampus. To determine the colocalization of P2X7R with astrocytes and microglia and its association with tau and amyloid pathology, triple staining was performed on 5 AD and 4 NC hippocampal tissue slices. Antigen retrieval of the paraffin-embedded fixed human brain tissue sections was performed with citrate buffer. First, the sections were heated in buffer for seven minutes at 700 W, followed by 20 minutes at 280 W. Last, the sections were kept at room temperature without cover for 20 minutes. After blocking in 5% donkey serum in 1% Triton-PBS, the sections were incubated with primary antibodies against GFAP, anti-calcium-binding adapter 1 (Iba1), anti-phospho-Tau (AT-8), anti-amyloid-beta (6E10, 4G8), or P2X7R overnight at 4°C with mild shaking (**Supplemental Table 1**) (44, 48).

For mouse brain sections, sagittal brain sections (40 μm) were cryosectioned. The sections were first washed in PBS 3×10 minutes, followed by antigen retrieval for 20 minutes in citrate buffer at room temperature. The sections were permeabilized and blocked in 5% normal donkey or goat serum and 1% Triton-PBS for one hour at room temperature with mild shaking. Free-floating mouse brain tissue sections were incubated with primary antibodies against GFAP, Iba1, AT-8, 6E10, CD68, and P2X7R overnight at 4°C with mild shaking (**Supplemental Table 1**) (31, 32, 69). The next day, the sections were washed with PBS for 2×20 minutes and incubated with a suitable secondary antibody for 2 hours at room temperature. The sections were incubated for 15 minutes in DAPI, washed two times for 10 minutes with PBS, and mounted with VECTASHIELD Vibrance Antifade Mounting Media (Vector Laboratories, Z J0215).

The H&E-, immunohistochemistry- and immunofluorescence-stained brain sections were imaged at ×20 magnification using an Axio Oberver Z1 slide scanner (Zeiss, Germany) with the same acquisition settings for all brain slices. The immunofluorescence-stained brain sections were also imaged at ×10 and ×63 magnification using a Leica SP8 confocal microscope (Leica, Germany).

### Image analysis

For the human hippocampus, manual delineation of the adult human brain based on the Allen atlas (21) was performed using Qupath and ImageJ (NIH, U.S.A.). CA1, CA2/3, the dentate gyrus (DG), the subiculum, the entorhinal cortex (EC), and the parahippocampal cortex (PHC) were drawn. The mean intensity was calibrated with the background intensity of each scan.

To quantify the regional fluorescent signal of GFAP, Iba1 and P2X7R based on confocal images, z-project with maximum intensity was applied followed by the HiLo (high low) look-up table in ImageJ (NIH, U.S.A.). Three background regions of interest (ROI) were manually drawn and their average background and average background per pixel acquired from raw integrated densities (RawIntDen) were calculated. To determine the intensity of the fluorescent signal, the RawIntDen of a large image ROI minus the average background per pixel multiplied by the area of the large ROI was quantified.

### Statistics

GraphPad Prism (GraphPad v10) and Python was used for statistical analysis. The normality of the data in the AD and NC cohorts was analysed by using the Shapiro□Wilk test. The nonparametric Mann□Whitney test was used to compare multiple groups. Braak stages (stage 0, I, II, III-IV, and V-VI), comparisons of Aβ levels (O-C converted to 0-4), and comparisons between males and females and between different hippocampal subregions. Nonparametric Spearman’s rank correlation analysis was performed between age, Braak stage, amyloid level and P2X7R level.

## Results

### Demographics and pathological description

The ages of the individuals in the NC group (82.2 ± 7.4 [67-99]) were comparable to those in the AD group (82.0 ± 8.4 [64-98]) and followed a normal distribution (Shapiro□Wilk test, p = 0.4029 and p = 0.5376, respectively). A greater percentage of females was noted in the AD group (88.2%) than in the NC group (61.3%). The prevalence of apolipoprotein E (APOE) ε4 carriers was significantly greater in the AD group (58.8%) than in the NC group (12.9%). No abnormalities were observed in the brain tissue slices from the AD patients or NCs based on H&E staining (**SFig. 1**). All the AD patients had an amyloid level of B or C and a varying Braak stage (II-VI). There are NCs that exhibit moderate amyloid-beta or tau pathology in the brain based on the amyloid and Braak stage scores (2). Braak stage and amyloid level information are described in **Table 1**.

### Reactive astrocytes and activated microglia in the hippocampus of AD patients and controls

To characterize the microglia and astrocytes in the AD and control brains, we first conducted immunohistochemical staining for Iba1 and GFAP in the hippocampi of 34 AD patients and 30 NCs. According to the Iba1 staining results, the Iba-1-positive microglia in the hippocampus of the AD patients appeared to have an increased volume and more pronounced ramifications than did those in the NCs (**Fig. 1a-j**).

**Fig. 1.**
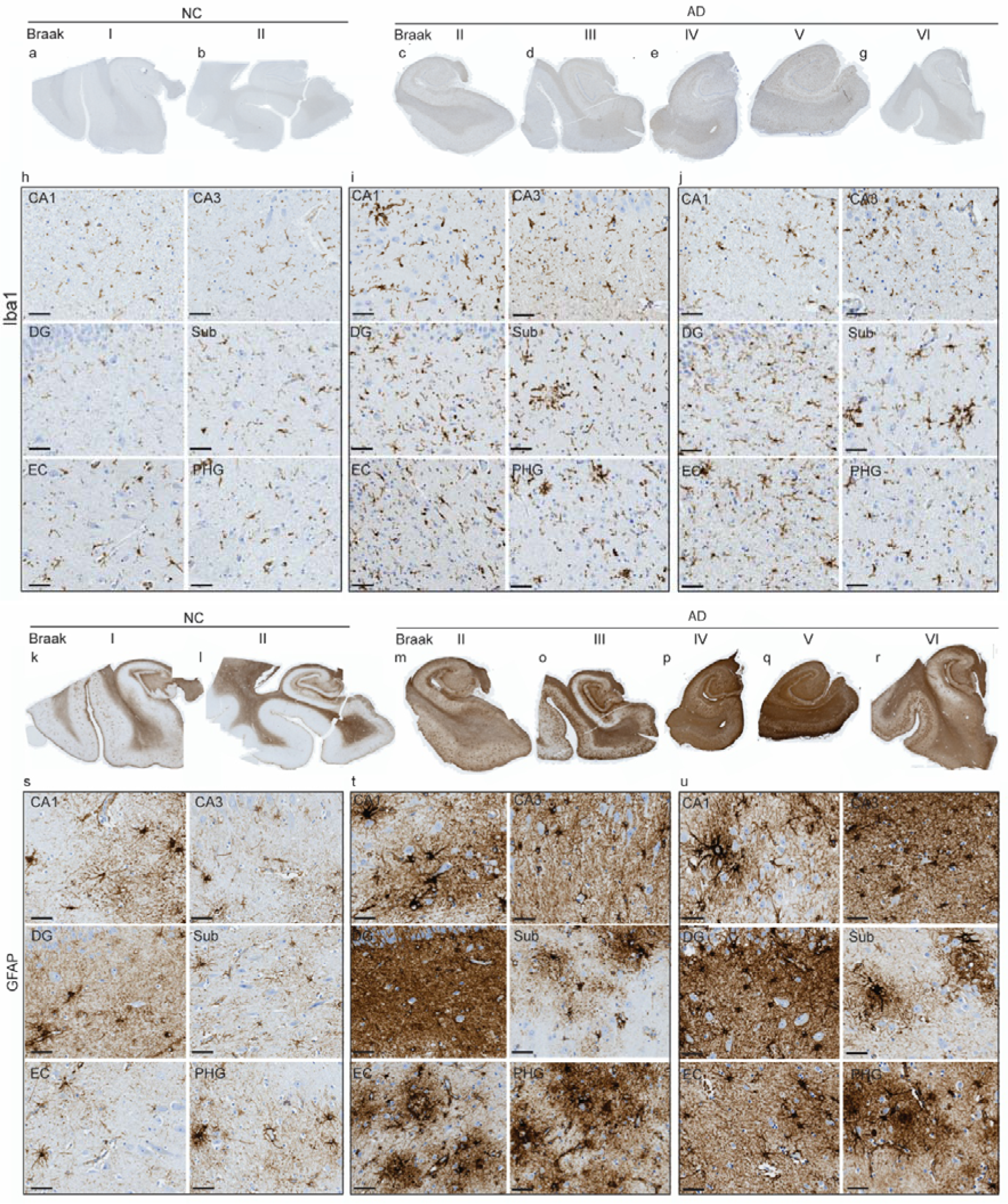
Iba-1 and GFAP staining of microglia and astrocytes in the hippocampi of non-demented control cases and AD patients. (**a-j**) Representative Iba1 (brown) and (**k-t**) GFAP (brown) immunohistochemical staining of the hippocampus of nondemented control cases (NC #04-015, #98-182) and AD patients (#99-127, #97-027, #99-139, #94-29, #93-12) at different Braak stages. Zoomed-in views of Iba1 (**h-j**) and GFAP (**r-t**) in the CA1-3 region, dentate gyrus (DG), Subiculum (Sub), entorhinal cortex (EC) and parahippocampal cortex (PHC) from (NC #04-015, AD #99-127, #93-012). Scale = 50 μm (h-j, r-t).

The cell bodies in the hippocampus of the AD patients were clearly observed via GFAP staining. The astrocyte processes became thicker in the AD hippocampus (**Fig. 1k-u**). The number of GFAP-positive astrocytes in the DG did not differ between AD patients and NCs. However, compared with those in NCs, more GFAP-positive astrocytes were observed in the CA1 region and subiculum of AD patients. Morphologically, GFAP immunohistochemical staining revealed that the astrocytes became denser astrocytes with reduced volume in the AD patients compared with the NCs. Hence, GFAP signal is condensed in the soma of cells from AD patient with reactive astrogliosis, whereas in NC, there is more pronounced axonal expression of GFAP. Similarly, Iba1-positive and activated microglia become larger, with increased ramifications upon disease progression.

### The P2X7R level in the hippocampus was greater in AD patients than in NC cases

Next, we quantified P2X7R expression in subfields of the hippocampus in the same cohort of 34 AD patients and 30 HCs by using immunofluorescence staining. The histogram of P2X7R level in the hippocampus of NC subjects and AD patients was shown in **SFig. 2**. The P2X7R level was quite even in the hippocampus and was slightly greater in the entorhinal cortex (EC), CA3 than in the CA1, subiculum (Sub), and dentate gyrus (DG) in both the AD and NC groups (**Fig. 2, 3a**). Given that not all the tissue blocks contained parahippocampal cortex, we did not include it in the data analysis.

**Fig. 2.**
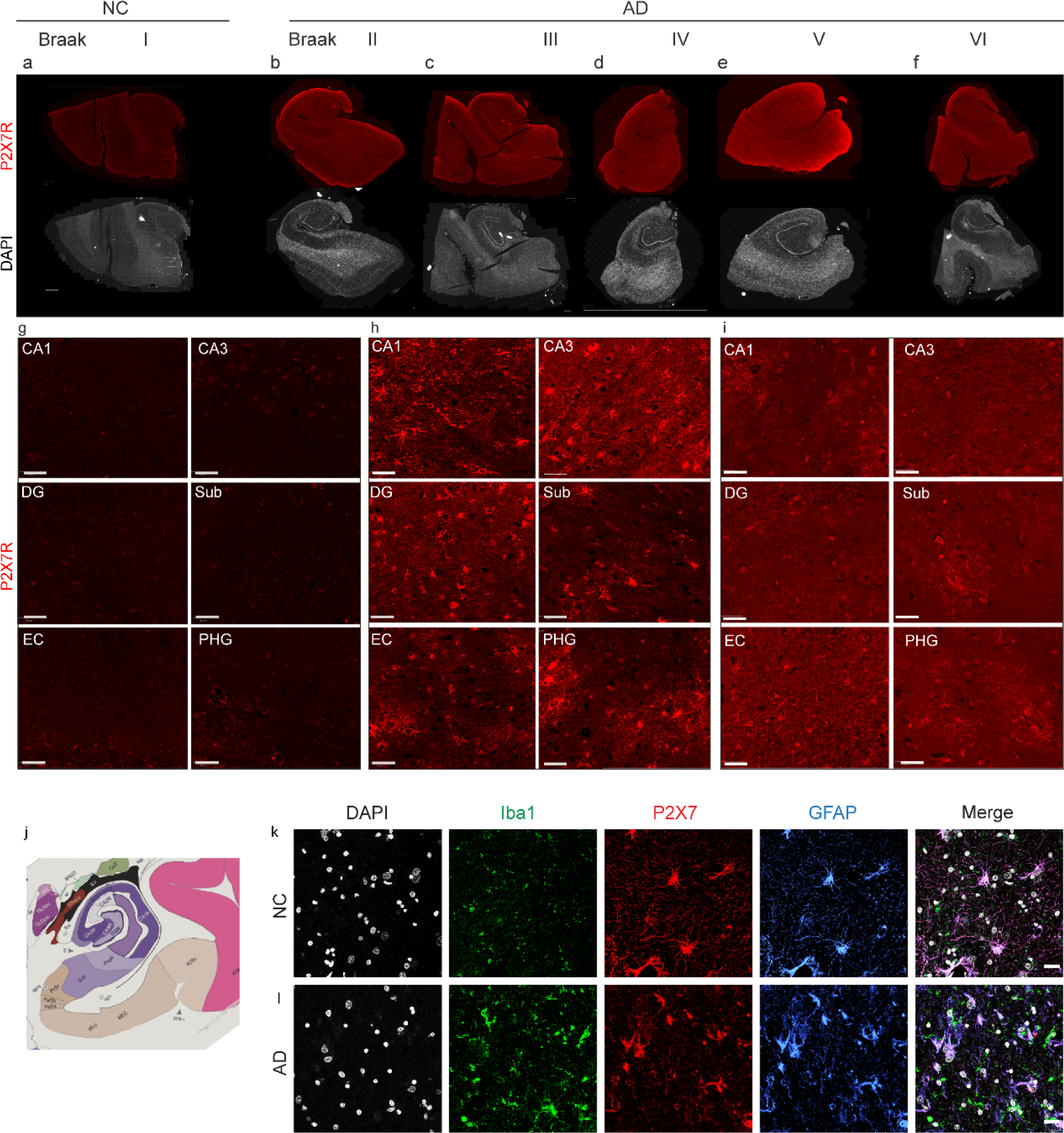
P2X7R staining in the hippocampi of NC and AD patients. (**a-i**) Representative P2X7R (red) immunofluorescence staining of the hippocampi of NC (#04-015, #98-182) and AD patients (#99-127, #97-027, #99-139, #94-29, and #93-12) at different Braak stages. (g-i) Zoomed-in views of P2X7R (red) (NC #04-015, AD #99-127, #93-012). CA1-3, dentate gyrus (DG), subiculum (Sub), entorhinal cortex (EC) and parahippocampal cortex (PHC). (j) Representative image from the Allen Brain Atlas. (k, l) P2X7Rs were colocalized with astrocytes and microglia in the hippocampus (DG). NC: #05-044, AD patient #97-051. Iba1 (green)/P2X7R (red)/GFAP (blue). Nuclei were counterstained with DAPI (white). Scale = 50 micron (g-i), 20 micron (k-l).

We found a significantly greater level of P2X7R (approximately 2×) in all the hippocampal subfields analysed in the AD patients than in the HCs (**Fig. 2a-j; 3**, CA1, p = 0.0004; CA3, p = 0.0004; DG, p = 0.0006; Sub, p = 0.0004; EC, p = 0.0002). Next, we performed confocal microscopy to assess the location of P2X7Rs on astrocytes and microglia in the hippocampus. P2X7R was observed both on Iba1-positve microglia as well as GFAP-positive astrocytes (**Fig. 2-4**). In the dentate gyrus, where astrocytes are highly expressed, more astrocytic P2X7Rs than microglial P2X7Rs were observed. The integrated intensity of GFAP (astrocytes) and Iba1 (microglia) did not differ in the hippocampus of the AD and NC group.

To understand whether the age and sex of the patients affect hippocampal P2X7R levels, correlation and group analyses were performed (**Fig. 3**). No sex difference in hippocampal P2X7R expression was detected between the AD group and the NC group (**Fig. 3b**). No correlation was observed between age and P2X7R expression in any of the hippocampal subfields in the AD or NC groups (**Fig. 3c**). APOE is a lipid transporter produced predominantly by astrocytes in the brain and plays an important role in modulating microglial immunometabolism (22). APOE ε4 is a genetic risk factor for sporadic AD.. Therefore, we assessed whether there was a difference between APOE ε4 carriers and noncarriers in terms of P2X7R levels. No difference was detected in the NC or AD groups between APOE ε4 carriers and noncarriers (**Fig. 3d**).

**Fig 3.**
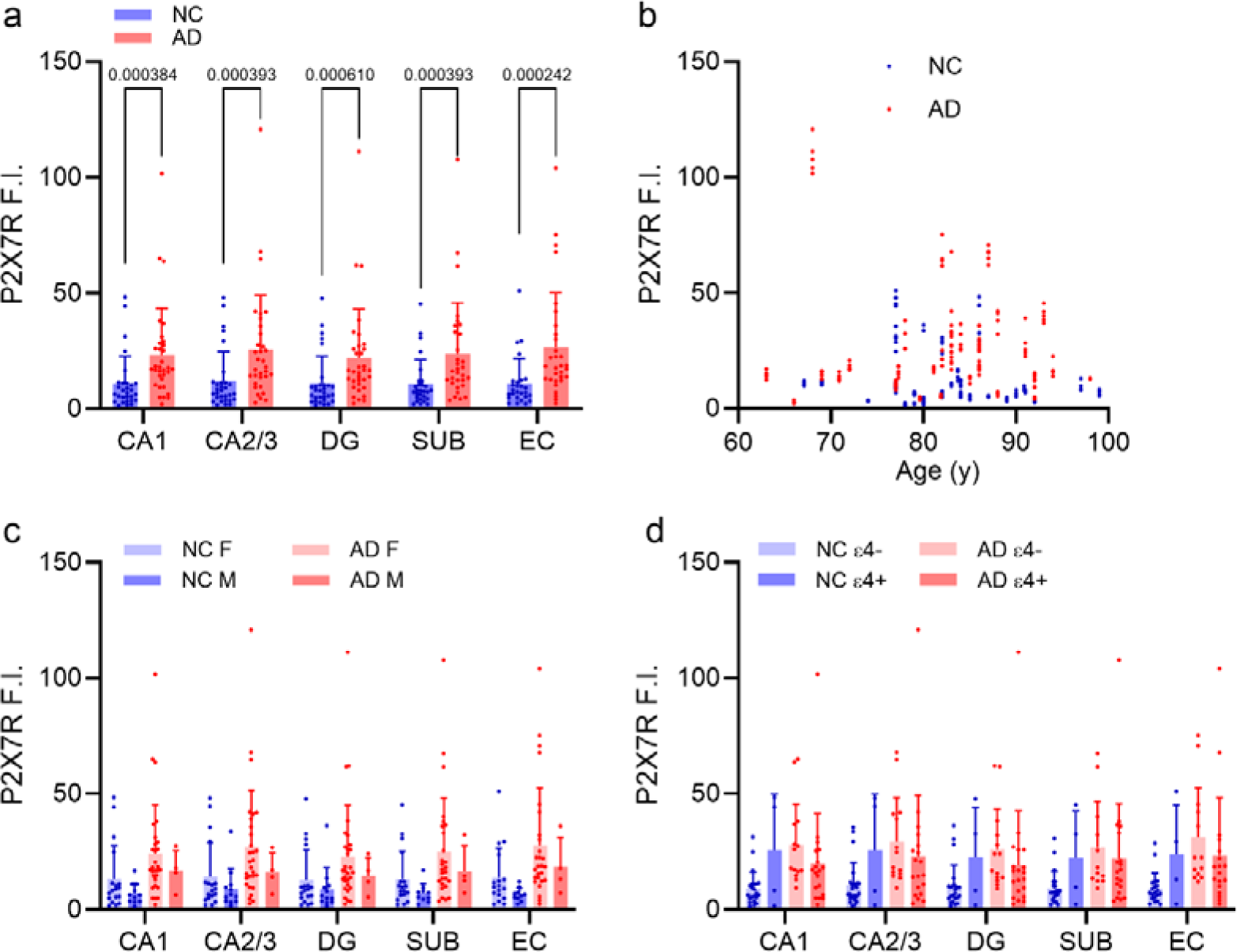
Increased P2X7Rs in the hippocampus of AD patients compared to NCs. (a) Distribution of P2X7Rs in the hippocampus subfields of AD patients (n= 34) and NCs (n= 30). (b) P2X7R expression did not correlate with age in the AD or NC group. (c) There was no difference in P2X7R expression between male and female NCs (n= 12, n= 18) or between male and female AD patients (n= 4, n= 30). (d) There was no difference in P2X7R levels between NC APOE ε4 carriers (n= 4) and noncarriers (n= 23) or between AD APOE ε4 carriers (n= 20) and noncarriers (n= 14). Dentate gyrus (DG), subiculum (SUB), and entorhinal cortex (EC). Nonparametric Mann□Whitney test.

### Associations between P2X7Rs and A**β** deposits in the hippocampi of AD patients

Next, we assessed the association between P2X7R and Aβ accumulation. Slidescanner and confocal microscopy were performed on P2X7R/6E10/Iba1-stained hippocampal tissue slices from 5 AD patients and 3 NCs. Amyloid plaques were detected in different subfields of the hippocampus of AD patients (**Fig. 4h, i**). The immunofluorescence reactivity of P2X7R increased around and inside the Aβ plaques, and close to cerebral amyloid angiopathy (**Figs. 4c-e**). Nonparametric Spearman’s rank correlation analysis revealed that the P2X7R level was positively correlated with the amyloid score in the AD and NC brains (p= 0.0129, r= 0.3274 in the CA1, p=0.0342, r=0.2786 in the CA3; p=0.0349, r=0.2752 in the DG; p=0.0019, r=0.4141 in the EC; p = 0.0060, r= 0.3657 in the Sub) (**Fig. 4j-n**).

**Fig. 4.**
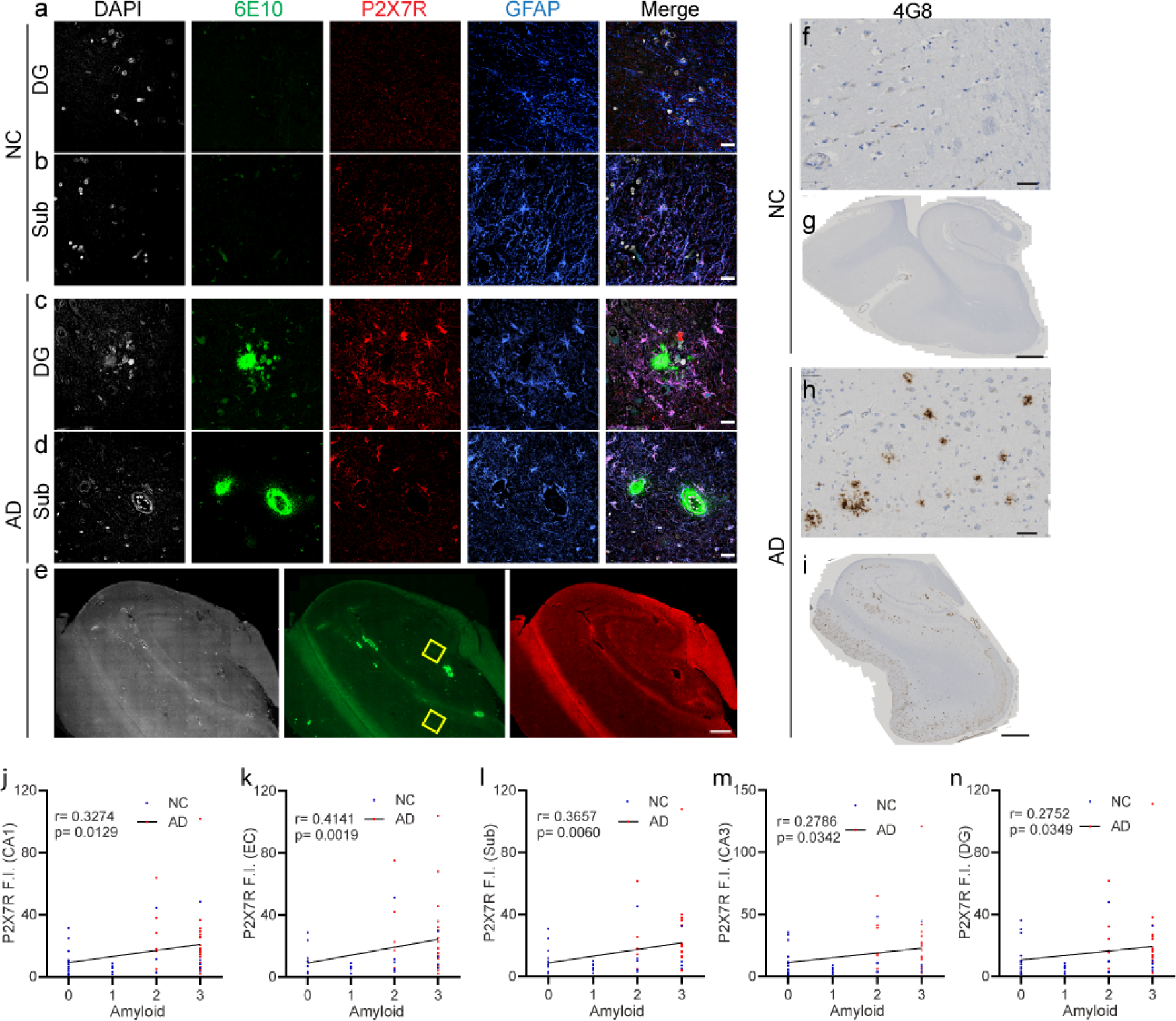
Increased P2X7R with amyloid-beta deposits in the hippocampus in AD patients. (**a-e**) Immunofluorescence staining on hippocampus sections from NC and AD patients using 6E10 (green)/P2X7R (red)/GFAP (blue). (**a-d**) Zoomed-in view of the dentate gyrus (DG) and subiculum (Sub). Increased P2X7R was detected surrounding parenchymal amyloid-beta plaques as well as cerebral amyloid angiopathy in the vessel wall. The yellow square in e indicates the location of the zoom in (c, d). Nuclei were counterstained with DAPI (white). (f-i) 4G8 Immunochemical staining for amyloid-beta on hippocampus sections from NC and AD patients Scale bar =20 μm (a-d), 50 μm (f, h), 800 μm (e), 2 mm (g, i). NC #08-010, AD #92-088. Nonparametric Spearman’s rank analysis of amyloid level (CERAD amyloid score O-C converted to 0-3 scale) and P2X7R expression in AD patients (n=30) and NCs (n=30).

### Associations between P2X7R and tau inclusions in the hippocampi of AD patients

Next, we assessed the associations of the accumulation of phosphorylated tau species and the Braak stage with P2X7R levels in the AD brain. Here, we used an AT-8 antibody (phospho-Ser202/phospho-Thr205 tau) to characterize the distribution of tau in the hippocampus (8). The pathology of tau in different stages of sporadic AD (9) and its spread in subregions of the hippocampus (from the EC via the prosubiculum to the dentate fascia) have been well characterized in the postmortem AD brain (10, 39, 62). We performed slidescan and confocal microscopy on P2X7R/AT-8/GFAP-stained hippocampi from 5 AD patients and 3 NC individuals. P2X7R increased around and colocalized with AT-8-positive tau inclusions (**Fig. 5d**). Increased GFAP-positive astrocytes around tau inclusions were also observed (**Fig. 5**). Nonparametric Spearman’s rank correlation analysis revealed that the P2X7R level was positively correlated with the Braak stage in the AD and NC brains (p= 0.0036, r=0.3796 in the EC; p = 0.0093, r=0.3386 in the Sub; p = 0.0097, r=0.3288 in the CA1). Given that the Braak staging does not indicate the tau level in the hippocampus, we performed phospho-Tau staining (using AT8 antibody) on the hippocampus slices of the AD patients and NCs.

**Fig. 5.**
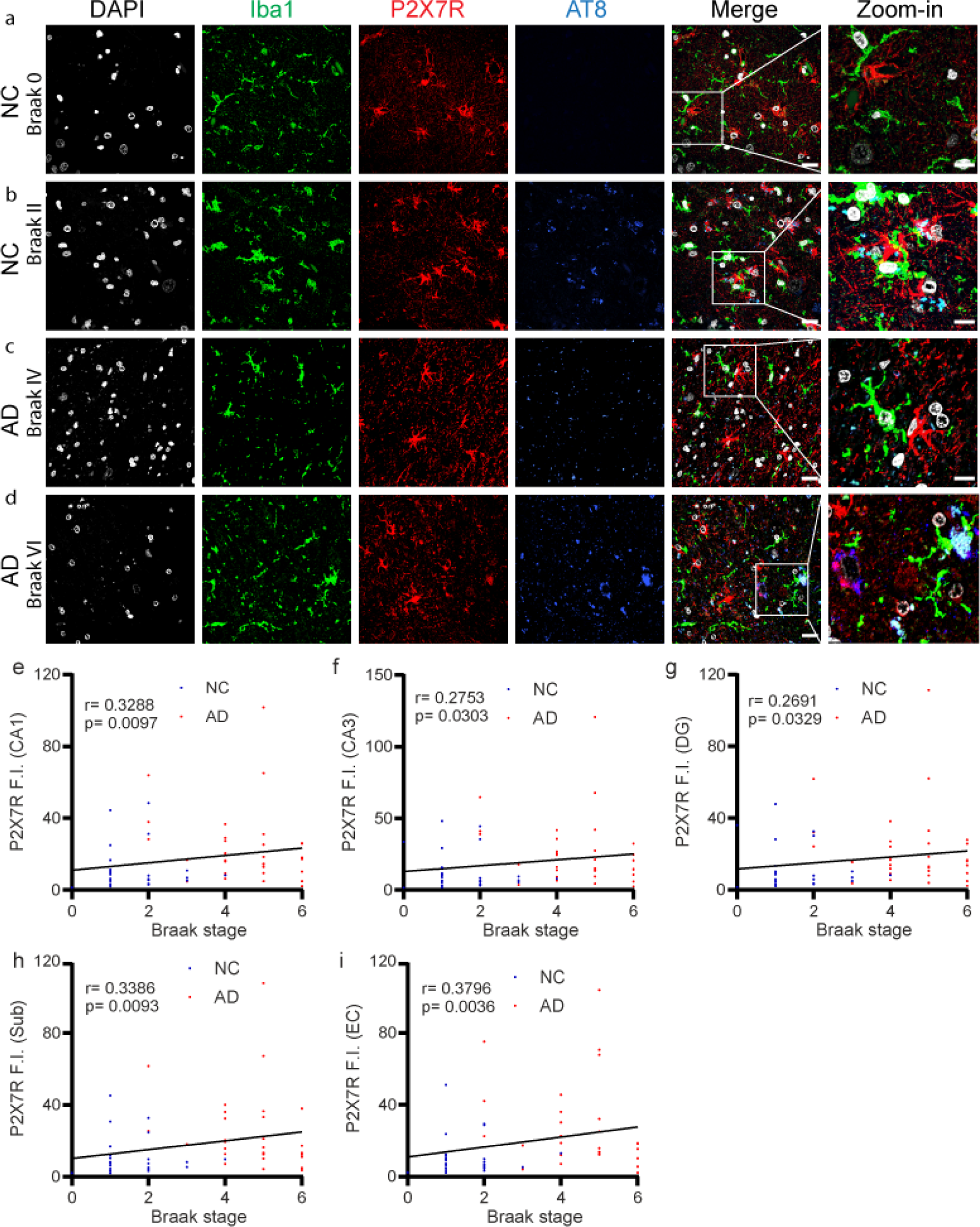
P2X7R colocalized phospho-Tau inclusion in the hippocampus of AD patients and correlated positively with Braak stage. (**a-d**) Hippocampal sections from the NC group (#99-249, #97-163); AD group (#92-088, #97-051) were stained for Iba1 (green)/P2X7R (red)/AT-8 (blue) to determine the association between P2X7R and phospho-Tau. Colocalization of tau and P2X7R and iba1+ activated microglia was observed. Nuclei were counterstained with DAPI (white). Scale bar = 10 μm. (**e-i**) Nonparametric Spearman’s rank analysis of Braak stage and P2X7R expression in AD patients (n=34) and NC (n=30).

### Associations between P2X7Rs and tau inclusions in the caudate/putamen of CBD and PSP patients

To further assess whether there were tau-related alterations in the levels of P2X7R, triple staining for microglia, P2X7R, and phospho-Tau was performed on the caudate/putamen slice from one CBD patient and on the putamen/insula slice from one PSP patient. Colocalization between AT-8-positive phospho-tau inclusions and P2X7Rs was observed in the caudate putamen slice from the CBD patient and the putamen insula slice from the PSP patient (**Fig. 6d, e**).

**Fig. 6.**
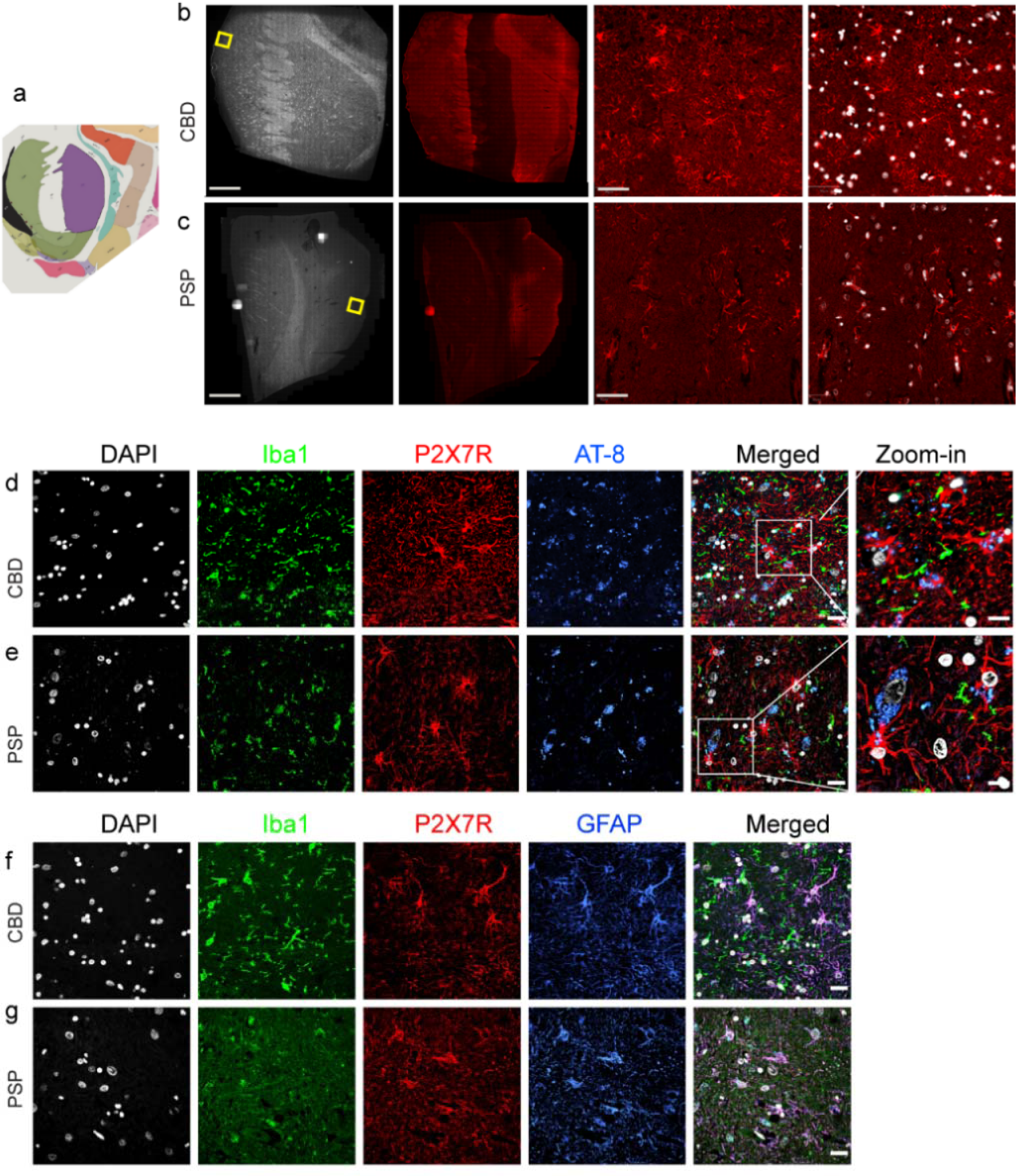
P2X7Rs in the putamen of the CBD and PSP brain. (**a**) Representative image from the Allen Brain Atlas. (**b-c**) Overview of DAPI/P2X7R (red) staining of cau/put sections from CBD patients and put/ins sections from PSP patients. The yellow square indicates the zoom-in image in columns 3 and 4. (**d, e**) Iba1 (green)/P2X7R (red)/AT-8 (blue) staining showing the association between P2X7R and phospho-Tau in the caudate/putamen section of a CBD patient and in the put/ins section of a PSP patient. (**f, g**) Iba1 (green)/P2X7R (red)/GFAP (blue) staining showing the localization of P2X7R on astrocytes and microglia. Nuclei were counterstained with DAPI (white). Scale bar = 2 mm (1, 2 column of a, b), 50 μm (3, 4 column of a, b); 20 μm (d-g column 1-5) and 10 μm (d, e column 6). CBD, corticobasal degeneration; PSP, progressive supranuclear palsy.

### Increased P2X7R in the hippocampus of 27-month-old 3×Tg mice

Next, we assessed P2X7R expression in sagittal brain tissue slices from wild-type and 27-month-old 3×Tg mice. Increased levels of P2X7R and CD68, which are indicative of microglial activation, were found in the subiculum of 3xTg mice compared to wild-type mice (**Fig. 7a-c**). 3×Tg mice exhibited tau and amyloid pathology in both the cortex and hippocampus (especially the subiculum) at 27 months of age (**Fig. 7f-h**). A greater level of P2X7R was especially observed surrounding the plaque, which was observed in the DAPI channel (**Fig. 7b**). We performed the staining for translocator protein (TSPO) in the same mouse brains. Higher TSPO signals are observed also in the white matter, where as the distribution of P2X7R is more in the grey matter of the mouse brain (**SFig. 3**).

**Fig. 7.**
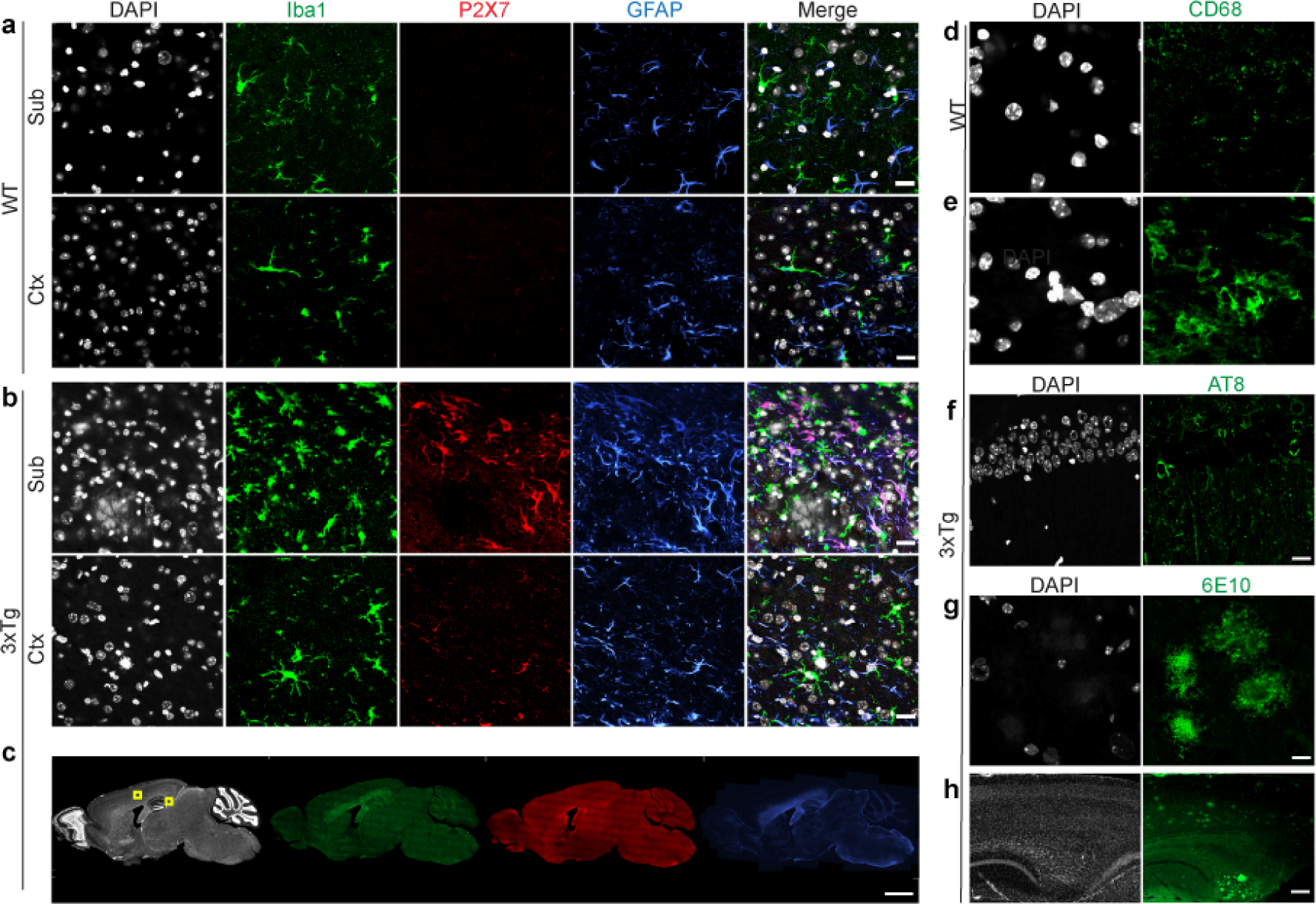
Immunofluorescence staining indicated increased levels of P2X7R, GFAP and Iba1, with the presence of phospho-Tau and amyloid deposits in the cortex and hippocampus of 27-month-old 3×Tg mouse brains. (**a-c**) Sagittal brain tissue sections from 27-month-old WT and 3×Tg mice were stained for Iba1 (green)/P2X7R (red)/GFAP (blue) in the Ctx (Layer 3/4) and Subiculum of the Hippocampus. (**d, e**) Images of CD68 (green) staining in the subiculum of WT and 3×Tg mouse brains. (**f, g, h**) AT-8 (green) and 6E10 (green) staining showing the presence of phospho-Tau and amyloid plaques in the subiculum of 27-month-old 3×Tg mice. Nuclei were counterstained with DAPI (gray). Scale bar = 20 μm (a, b, d-g). 200 μm (h), 1 mm (c). Yellow squares indicate the locations of the zoomed-in views.

## Discussion

Here, we showed that the hippocampal level of P2X7R was greater in AD patients than in NCs, and correlated with amyloid pathology and tau pathology indicated by Braak stage. We found no age-related, APOE ε4 allele-related or sex-related differences in the hippocampal P2X7R levels between the AD and NC groups.

Both increase and no difference have been reported earlier in terms of the P2X7R level in AD brain tissue compared to NCs. Similar to our observation, the age or sex related difference was also not reported in a recent study on P2X7R using temporal cortex tissue from 35 AD and 25 NCs (4). Our finding is in line with a previous study showing an increased P2X7 protein levels by Western blotting and P2X7 mRNA levels in the hippocampal homogenates from 9 AD and 6 NCs (17). Other studies have investigated the P2X7R changes in the cortex: an increased expression and activation of P2X7R have been reported in three postmortem studies on the temporal cortex tissue of 5 AD vs 4 NC, 5 FTLD, 10 PSP, 5 CBD vs. 4 NC, and microglia from 6 AD vs 8 NC (13, 42, 45). A recent study showed that P2X7R levels in the temporal cortex tissue were higher only in 16 AD of Braak V-VI compared to 25 NC of Braak 0-II, but not between 19 AD of Braak III-IV compared to NC group (4). However, another study showed immunohistochemical evidence of no difference in the levels of P2X7R, P2X4R and CaMKK2 in pyramidal neurons of the frontal cortex in AD patients and control case (24). An earlier autoradiography study using P2X7R tracer [^11^C]SMW139 showed comparable P2X7R levels in temporal cortex tissue from AD patients and nondemented controls (NCs) (29). A recent study showed a significant increase in plasma P2X7R levels in AD patients compared to those in control individuals (sensitivity 90.0%, specificity 50.22%) or patients with mild cognitive impairment (1). We found that the number of GFAP+ astrocytes and iba1+ microglia was not increased in the CA1, CA3 region of the hippocampus despite the increase in the level of P2X7R. Earlier study have performed Stereological layer-specific analysis based on the IHC and proteomics on the EC tissue from AD and NC (3). The observation of no change in Iba1+microglia is in line with earlier study where No significant differences were observed in the total number of identified microglia between controls and AD patients, neither in the grey or white matter (33). The immunohistochemical staining stereological analysis is considered as most accurate quantitative method for astrocyte and microglia analysis. However this method is time-consuming and laborious, therefore we analysed here using immunofluorescence staining based quantification.

We found that the level of P2X7R was positively associated with amyloid pathology and tau pathology indicated by Braak stage in the hippocampi of AD patients and NC. Increased P2X7R was observed surrounding amyloid plaques and tau inclusions in the AD brain, around the tau inclusion in the caudate/putamen of CBD and PSP patients. Moreover, in aged 3×Tg mice, increased P2X7R was observed in the subiculum with amyloid pathology based on the staining results. We recently showed an increased uptake of the P2X7R tracer [^18^F]GSK1482160 in vivo in the brains of 7-month-old rTg4510 mice with tauopathy, which correlated with [^18^F]APN-1607-measured tau accumulation in the hippocampus (37, 69). This is in line with a recent study using in-situ hybridisation-immunohistochemistry showing that P2RX7 mRNA localises to GFAP+ astrocytes and CD68+ microglia surrounding Aβ plaques in AD brain (4). Studies have shown that the use of P2X7R antagonists reduces cell death by altering the balance of tau phosphorylation inside and outside cells by reducing tissue-nonspecific alkaline phosphatase (TNAP) expression (17). These results suggest a close link between P2X7R and amyloid and tau pathologies.

The association between microglia, astrocyte with AD pathologies has not been fully understood. Aβ and tau pathologies have been shown to exert distinct influences and induce disease stage-specific microglial subtypes (25, 35). Microglial activation measured by PET using tracer for translocator protein is found associated with AD amyloid and tau pathology (58). While another study indicated that microglial activation protects against the accumulation of tau in individuals without dementia (51). In both AD, and animal model of amyloid and tauopathy, PET evidence of microglia and astrocytosis are observed (28, 57, 70).

Hippocampal GFAP-positive astrocytes have also been shown to respond to Aβ and tau pathologies (16). In turn, severe neuroinflammation does not intensify neurofibrillary degeneration in the human brain (65). A recent postmortem analysis has also shown that tau oligomer-containing synapses are eliminated by microglia and astrocytes in AD (66). To date, there has been no in vivo imaging study of P2X7R in AD, as the optimal option P2X7R tracer suitable for central nervous system application is relatively limited. In vivo imaging of P2X7Rs with amyloid and tau imaging or fluid biomarkers will provide better insight into the potential associations between P2X7R and AD pathologies.

We found that similar increase of P2X7R was observed in the 27 month old 3×Tg mice compared to age-matched wild-type, especially in the subiculum where amyloid pathology locates. Elevated expression levels of P2X7Rs have been observed in different mouse models of AD amyloidosis such as APP/PS1 mice (40, 42), J20 mice (43), as well as in rats injected with Aβ_42_ (45), and in tauopathy rTg5410 (69) and THY-Tau22 mice (13).

There are several limitations in our study. First, there was a greater percentage of females in the AD group than in the NC group. Second, the gradient of P2X7Rs in the hippocampal head, body, and tail was not considered. Further in vivo PET using the P2X7R tracer in living patients will enable 3D analysis. Third, we did not have a clinical history of patients whose P2X7R level was correlated with that at postmortem. Moreover, for P2X7Rs, which are expressed on both astrocytes and microglia, further analysis of the extent to which astrocytic and microglial P2X7Rs are upregulated will provide further insights into the cellular source of P2X7. Here, we did not perform transcriptomic analysis, RNA analysis or profiling of P2X7R, P2X4R, or P2Y12R expression.

## Conclusions

In conclusion, we showed that hippocampal P2X7R expression was greater in AD patients than in NCs, associated with amyloid and tau pathology (Braak stage), and was not influenced by sex, age, or APOE ε4 allele. P2X7R has potential as a promising biomarker for neuroinflammation.

## Declaration

### Funding

RN received funding from the Swiss Center for Applied Human Toxicity (SCAHT-AP_22_01), Zurich Neuroscience Zentrum, and Helmut Horten Stiftung.

### Competing interests

CH and RMN are employees and shareholders of Neurimmune AG. Other authors declare no conflicts of interest.

## Acknowledgements

The authors acknowledge the Center for Microscopy and Image Analysis (ZMB), University of Zurich; Daniel Schuppli at Institute for Regenerative Medicine, University of Zurich.

## Author contributions

RN conceived and designed the study. CM performed staining and microscopy. CM, JW, and RN performed the data analysis. Postmortem human brain tissues were obtained from NBB. CM, RN wrote the first draft. All the authors contributed to the revision of the manuscript. All the authors have read and approved the final manuscript.

