## Supplementary figures for "Hippocampal purinergic P2X7 receptor level is increased in Alzheimer’s disease patients, and associated with amyloid and tau pathologies"

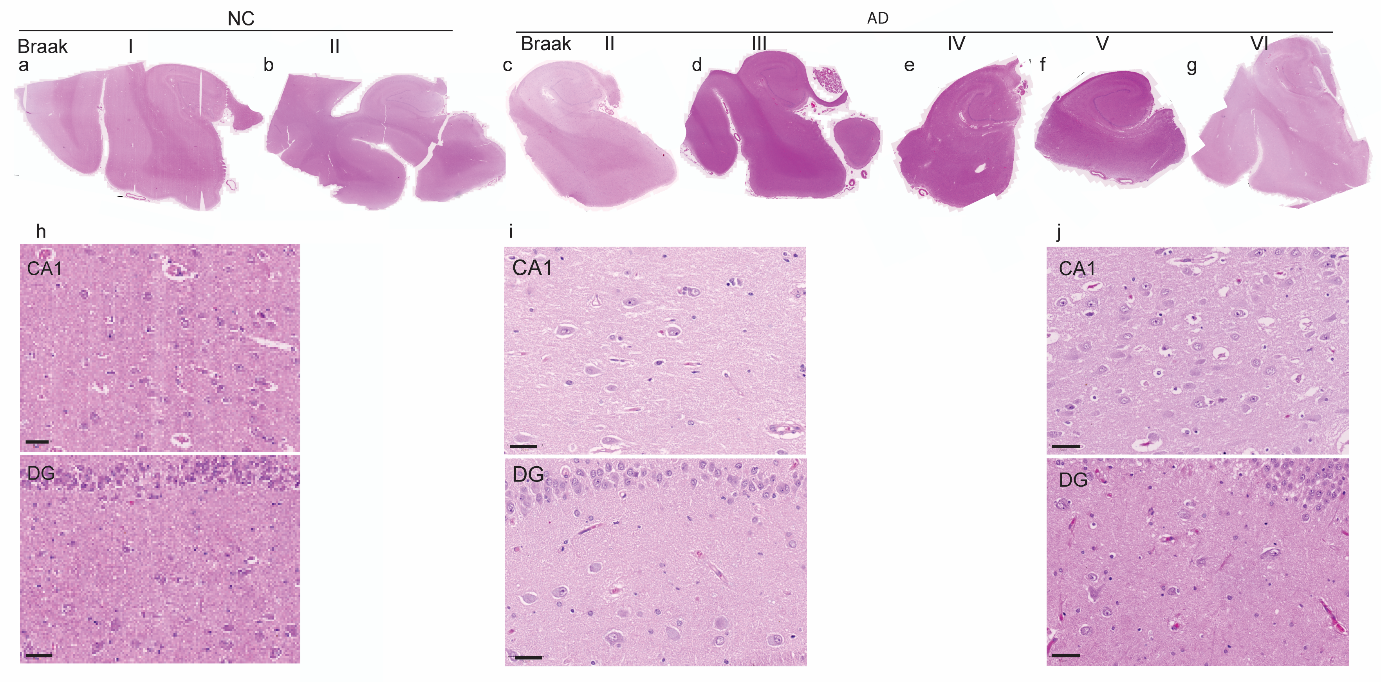

**SFig. 1 Hematoxylin and eosin staining of the hippocampi of control and AD patients. (a-j)** Representative H&E staining of the hippocampi of control patients (NC #04-015, #98-182) and AD patients (#93-12, #97-027, #94-29, #99-139, #99-127) at different Braak stages (shown in Fig 1). Zoom in views of CA1, dentate gyrus (DG).

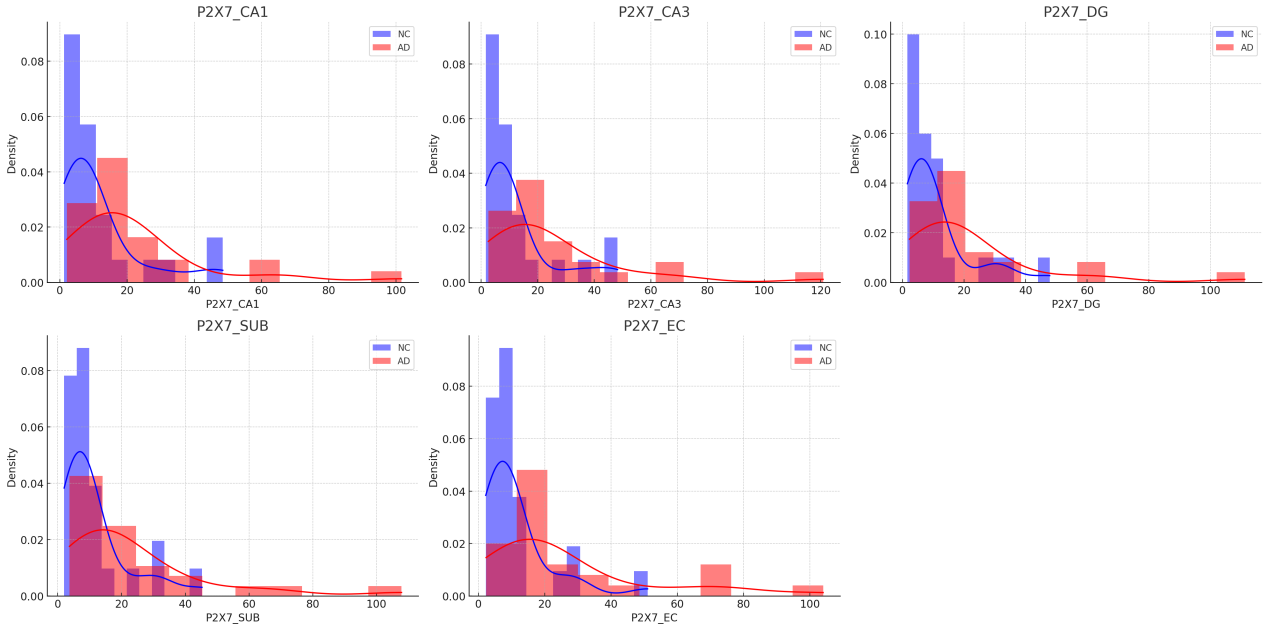
**SFig 2** **Histogram of the P2X7R level in the hippocampus of NC subjects and AD patients**.

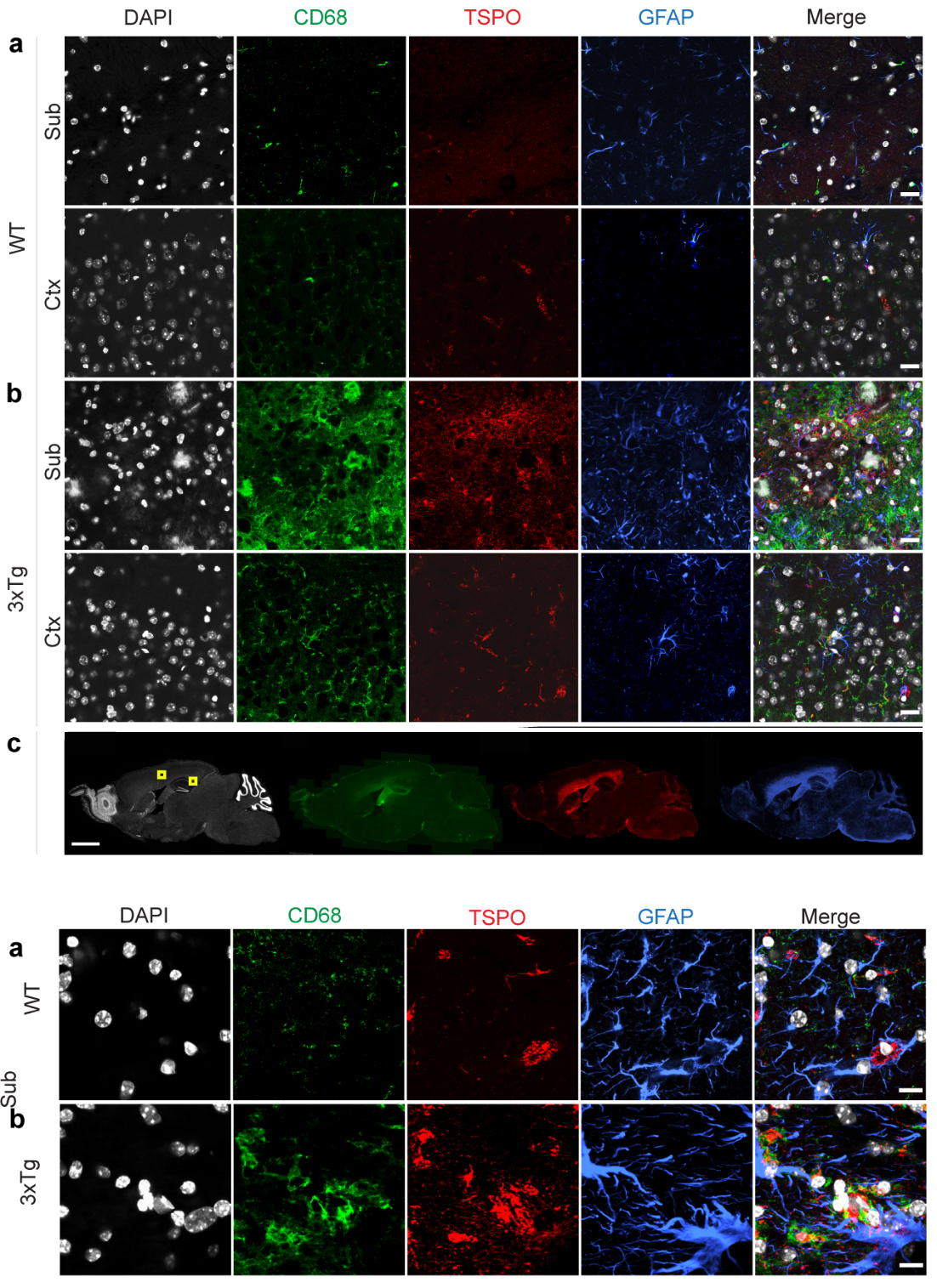

**SFig 3. Immunofluorescence staining indicated increased levels of TSPO, GFAP and CD68 in the cortex and hippocampus of 27-month-old 3×Tg mouse brains.** (**a-c**) Sagittal brain tissue sections from 27-month-old WT and 3×Tg mice were stained for CD68 (green)/TSPO (red)/GFAP (blue) in the Ctx (Layer 3/4) and Subiculum of the Hippocampus. Nuclei were counterstained with DAPI (gray). Scale bar = 20 μm (a, b). 1 mm (c). Yellow squares indicate the locations of the zoomed-in views.
