## Supplemental Table 1 for "Hippocampal purinergic P2X7 receptor level is increased in Alzheimer’s disease patients, and associated with amyloid and tau pathologies"

**Table S1 Antibodies and reagents used in the study**

| **Item** | **Catalog no** | **Dilution** | **Supplier** |
| --- | --- | --- | --- |
| Mouse phospho-Tau (Ser202, Thr205) monoclonal antibody (AT-8) | MN1020 | 1:1000 | Invitrogen |
| Mouse purified anti-β-Amyloid, 1-16 monoclonal antibody (6E10) | 803001 | 1:1000 | Biolegend |
| Mouse purified anti-β-Amyloid, 17-24 monoclonal antibody (4G8, IHC) | 800701 | 1:4000 | Biolegend |
| Guinea pig GFAP polyclonal antibody, IF | BP5082 | 1:1000 | OriGene |
| GFAP Monoclonal Antibody (6F2) IHC | MA1-35377 | 1:50 | Thermo Fisher |
| Rabbit ionized calcium-binding adapter molecule 1 (Iba1), IHC & IF | 019-19741 | 1:1000 | WAKO |
| Goat purinergic P2X7 receptor (P2X7R) | NBP1-37775 | 1:100 | Novus Biologicals |
| Rabbit recombinant Anti-CD68 antibody [EPR20545] | Ab213363 | 1:1000 | Abcam |
| Alexa fluor 488 donkey anti-rabbit IgG (H+L) | 711-545-152 | 1:250 | Jackson |
| Alexa fluor488 donkey anti-mouse IgG (H+L) | 715-545-151 | 1:500 | Jackson |
| Donkey anti-goat IgG (H+L) cross-adsorbed secondary antibody, Alexa fluor 546 | A-11056 | 1:50 | Invitrogen |
| Alexa fluor647 donkey anti-guinea pig IgG (H+L) | 706-605-148 | 1:250 | Jackson |
| Alexa fluor647 donkey anti-mouse IgG (H+L) | 715-605-151 | 1:250 | Jackson |
| DAPI (4',6-Diamidino-2-Phenylindole, Dihydrochloride) | D1306 | 1:1000 | Invitrogen |
